# Signal integration and information transfer in an allosterically regulated network

**DOI:** 10.1101/518514

**Authors:** Erin M. Shockley, Carol A. Rouzer, Lawrence J. Marnett, Eric J. Deeds, Carlos F. Lopez

## Abstract

A biological reaction network may serve multiple purposes, processing more than one input and impacting downstream processes via more than one output. These networks operate in a dynamic cellular environment in which the levels of network components may change within cells and across cells. Recent evidence suggests that protein concentration variability could explain cell fate decisions. However, systems with multiple inputs, multiple outputs, and changing input concentrations have not been studied in detail due to their complexity. Here, we take a systems biochemistry approach, combining physiochemical modeling and information theory, to investigate how cyclooxygenase-2 (COX-2) processes simultaneous input signals within a complex interaction network. We find that changes in input levels affect the amount of information transmitted by the network, as does the correlation between those inputs. This, and the allosteric regulation of COX-2 by its substrates, allows it to act as a signal integrator that is most sensitive to changes in relative input levels.

---

Many biological signaling networks process multiple inputs and yield multiple outputs. Examples of multiple-input multiple-output (MIMO) biochemical systems include the mitogen-activated protein kinase (MAPK) network, which can respond to numerous ligands and yield a range of outputs including proliferation and differentiation(Santos et al., 2007); the NF-*κ*B pathway, which triggers pro- and anti-inflammatory responses to a variety of ligands (Lawrence, 2009); and myriad metabolic networks, which respond to multiple substrates and allosteric regulators by producing energy and the building blocks of cellular components (Lorendeau et al., 2015). Recent work (Adlung et al., 2017; Spencer et al., 2009; Shi et al., 2016; Huang, 2009; Waite et al., 2016; Mitchell et al., 2018; Chen et al., 2012) has highlighted the fact that modulation of input concentrations in intracellular networks can yield markedly different outcomes. Despite this clear indication that MIMO systems are crucial to biological processes, few reports exist to date to explain how multiple inputs modulate reaction flux and information flow in a network to allow signal processing with a range of adaptive outputs.

To explore the properties of MIMO systems in biology, we chose to study the dynamics of cyclooxygenase-2 (COX-2), a key enzyme that controls the balance between pro- and anti-inflammatory signals in mammalian organisms. COX-2 lies at the interface of the eicosanoid and endocannabinoid signaling pathways (Alhouayek and Muccioli, 2014; Rouzer and Marnett, 2011) and is itself the target of the widely used nonsteroidal anti-inflammatory drugs (NSAIDs). Although COX-2 is a structural homodimer, it behaves as a heterodimer. One subunit in the dimer harbors the catalytically active site, while the other subunit contains an allosteric site that modulates the overall activity of the enzyme (Dong et al., 2013, 2011; Kulmacz and Lands, 1984). An array of substrates, inhibitors, and allosteric modulators can bind to, and thus compete for, either site, giving rise to highly complex reaction kinetics (Kudalkar et al., 2015; Kulmacz and Lands, 1985; Mitchener et al., 2015; Rimon et al., 2010; Yuan et al., 2009; Dong et al., 2016a). The various products from COX-2 activity drive multiple downstream pro- and anti-inflammatory processes that lead to diverse cellular fates including stress responses and apoptosis (Funk, 2001; Rouzer and Marnett, 2003; Smith et al., 2000).

It is clear that COX-2 orchestrates a complex interplay between a variety of substrates (the enzyme *inputs*), various allosteric regulators, and the concentration of downstream products (the enzyme *outputs*) that control processes such as inflammation (Funk, 2001; Rouzer and Marnett, 2003; Smith et al., 2000). Previously, most studies of COX-2 function have used simplified models based on Michaelis-Menten kinetics (Briggs and Haldane, 1925). Not surprisingly, these approaches have proved insufficient to capture the rich complexity of the COX-2 network of reactants, intermediates and products (Mitchener et al., 2015). We posit that a systems approach to understand COX-2 mechanism will improve inhibitor design to achieve desired outcomes in clinical settings.

COX-2 activity also represents an ideal model system to study the detailed dynamics of a biological MIMO system. As a single enzyme, it is sufficiently simple to allow for the construction, simulation and parameterization of a detailed systems biochemistry model that can capture all of the relevant transitions between intermediates and products. Nonetheless, it is sufficiently complex that it represents a non-trivial example of how multiple inputs lead to multiple outputs in a physiological context. We focus our study on the allosteric regulation network of COX-2 by two important substrates, arachidonic acid (AA) and 2-arachidonoylglycerol (2-AG), which generate unusual dynamics in the COX-2 network when both are present (Mitchener et al., 2015). Levels of AA and 2-AG also vary widely *in vivo* (Seibert et al., 1997; Monjazeb, 2006; Sugiura et al., 2006), and it is unclear how such variation would influence COX-2 signal processing.

In this work, we analyze the execution mechanism of a biochemical reaction network with multiple inputs. Our work explains how a MIMO system integrates information on the concentration and nature of its substrates to yield potentially different outputs. In previous work, we developed a detailed model of the COX-2 reaction network that comprises all possible biochemical enzyme states dictated by AA and 2-AG occupancy of the allosteric or active sites, and all the kinetic transitions between these states (Mitchener et al., 2015). The reaction rate parameters for the kinetic system were determined using a Bayesian inference methodology (Shockley et al., 2017) to fit the model to experimental data on COX-2 kinetics. This Bayesian approach produced an ensemble of model parameters that represent the uncertainty in the kinetic rates given the available data and restricts our analysis to plausible kinetic states of the network (Mitchener et al., 2015; Shockley et al., 2017). To explore the COX-2 MIMO signal processing mechanism, we first employed a graph-theoretic approach to enumerate all possible paths a substrate can take from reactant to product molecule. We found that changing the concentration of the inputs modulates not only the most dominant path that is taken by the system, but also the diversity of the paths the system employs. We also used an information-theoretic approach (Shannon, 1948) to understand the flow of information between network inputs, various intermediates, and the product outputs. This analysis reveals that competition between AA and 2-AG for the allosteric and active site generates highly complex concentration-dependence curves for COX-2 that are context-sensitive. In addition to providing insight into how COX-2 functions as a hub for the processing of inflammatory signals, our work suggests that our systems biochemistry framework provides useful information relevant to the study of other MIMO biological systems. This work also demonstrates that the extreme context-sensitivity of MIMO systems must be considered when attempting to modulate their behavior through targeted interventions.

## RESULTS

### A Mathematical Model of COX-2 Allostery and Catalysis

We built the COX-2 Reaction Model (CORM) (Fig. 1*B*) to understand how substrate-dependent allosteric regulation affects COX-2 catalytic rates (Mitchener et al., 2015). Here, we employ this model to study how multiple signals are processed in the context of a complex chemical reaction network, given a range of substrate concentrations and input correlations. Briefly, CORM encodes the reaction kinetics between COX-2 and two of its substrates: the fatty acid arachidonic acid (AA) and the endocannabinoid 2-arachidonoylglycerol (2-AG). Both AA and 2-AG can bind at the catalytic and/or allosteric site on COX-2 with different affinities. At the catalytic site, AA is turned over to prostaglandin (PG) while 2-AG produces prostaglandin-glycerol (PG-G). Binding of either molecule at the allosteric site modulates the rate of catalysis (Mitchener et al., 2015). Although CORM includes only two substrates, the MIMO nature of COX-2 kinetics results in a complex network (Fig. 1*B*). CORM has been calibrated to experimental data using PyDREAM, a Bayesian parameter inference framework, to obtain the probabilistic likelihood of parameters given experimental data, and information about the uncertainty in those parameter values (Shockley et al., 2017). CORM is encoded in Python using PySB, which provides a flexible tool to query the mixture of complexes present in the system at any time point given starting concentrations. Many of these complexes would be costly or impossible to measure experimentally. Employing the Python environment also facilitated the sophisticated analyses we present in this work (Lopez et al., 2013).

**Figure 1.**
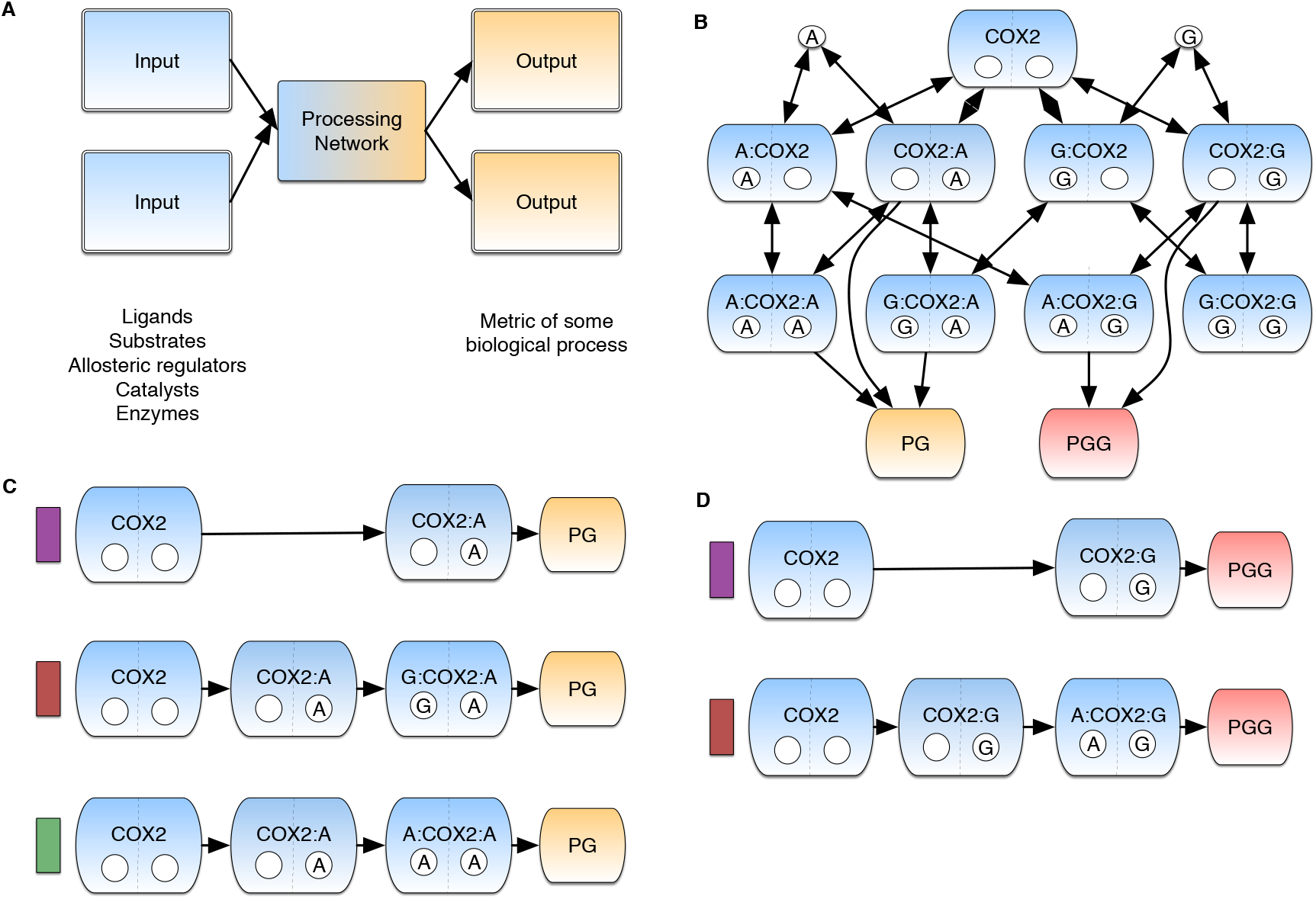
Network interactions within CORM. (*A*) The Multi-input Multi-output motif in a biological context. (*B*) The COX-2 Reaction Model (CORM) represents the network of interactions in the COX-2 system. The diagram depicts the possible biochemical states that the COX-2 enzyme (blue lozenges) can adopt through its allosteric (lower left circle) and catalytic (lower right circle) subunits, respectively. AA bound in either site is indicated with **A** and 2-AG with **G** within the circle. AA is turned over to produce prostaglandin (PG) and 2-AG is turned over to produce prostaglandin-glycerol (PG-G). Double-headed arrows indicate reversible reactions while single-headed arrows indicate irreversible reactions. Credible intervals for all fitted parameters are included in SI. (*C*) Dominant PG Production Paths in CORM. Colors correspond to path fluxes in Fig. 2*A*. (*D*) Dominant PG-G Production Paths in CORM. Colors correspond to path fluxes in Fig. 2*B*.

### Substrate-Dependent Reaction Fluxes in Signal Execution

We first explored the net flow of reaction flux through the network using a graph theoretic approach to calculate all possible paths between the unbound enzyme and each final product. Briefly, we evaluated the system of ordinary differential equations (ODEs) in CORM at time intervals to extract the integrated reaction flux at a given time point for each chemical reaction. We then built paths from product to reactant following the reactions with net forward flux. Finally, we calculated the total chemical flux that passed through a given path and used this as a measure of the probability of product formation via that path; a detailed description of this procedure is given in SI Methods and Fig. S1. All fluxes were calculated for the first ten seconds of catalysis after mixture with the substrates, a time chosen to match previous experimental work (Mitchener et al., 2015). Path flux distributions were calculated for an ensemble of calibrated parameter values to quantify path flux uncertainty arising from parameter uncertainty.

Our analysis indicates that there are six possible paths to produce PG (Fig. S2) and four possible paths to produce PG-G (Fig. S3) for all evaluated substrate concentration combinations. However, not all paths exhibit significant reaction flux during catalysis across all the concentrations. This occurs because paths in which binding of a species to the allosteric site precedes binding to the catalytic site are kinetically disfavored in CORM. As shown in Fig. 1*C* and 1*D*, three paths dominate PG production and two paths dominate PG-G production. The dominant PG-producing paths (Fig. 1*C*) include those with one or two intermediates, and the allosteric site empty or occupied by AA or 2-AG. Our results show that the dominant path is highly dependent on the substrate input concentrations. The presence of AA and 2-AG in the allosteric site enhances the production of PG (Mitchener et al., 2015). The dominant PG-G-producing paths include one or two intermediates (Fig. 1*D*) with the allosteric site empty or occupied by AA. The presence of AA in this site reduces the rate of PG-G production Mitchener et al. (2015). Similar to PG production, we also found that the flux through each dominant path for PG-G production is dependent on substrate concentration (Fig. 2).

**Figure 2.**
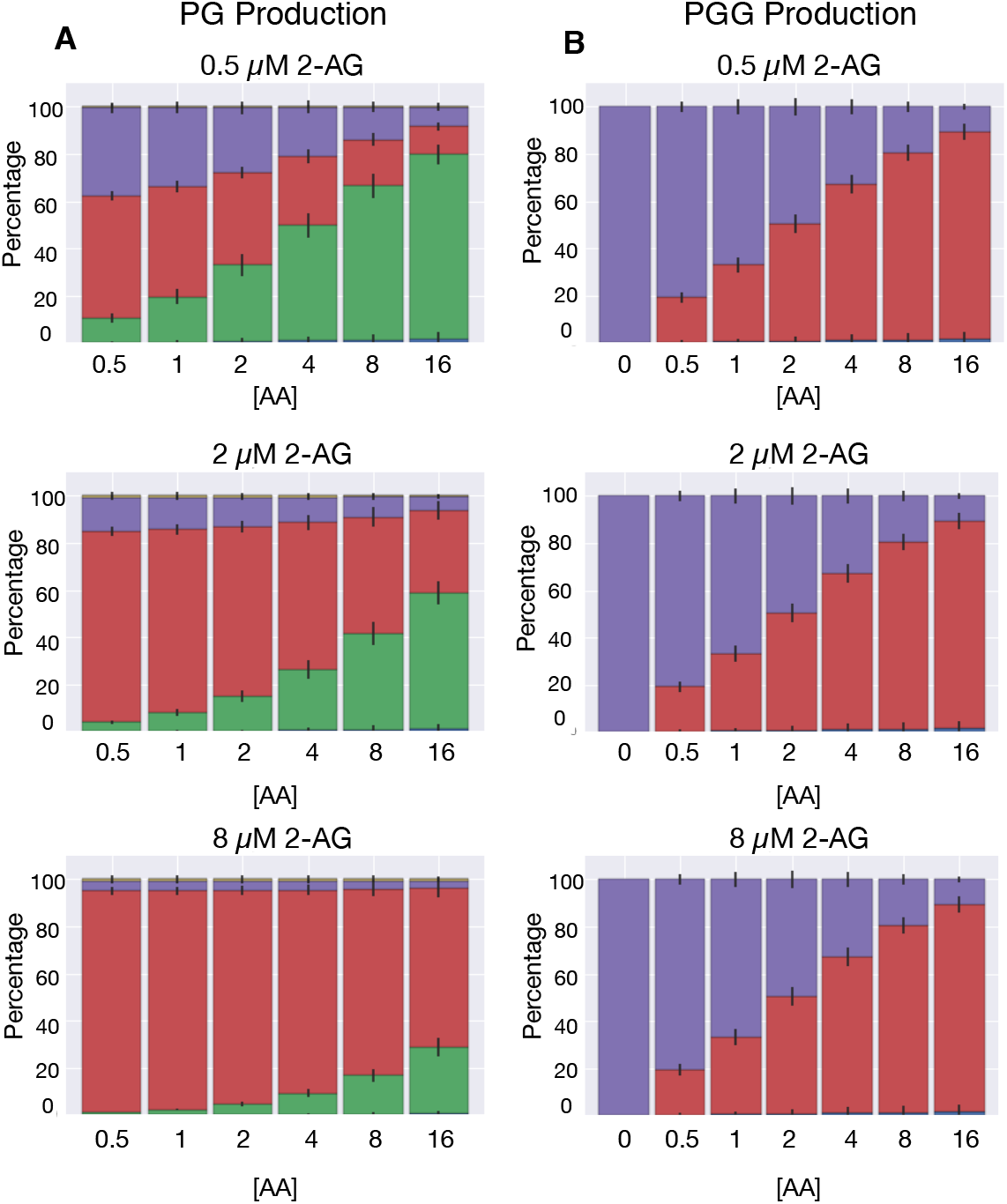
Concentration-dependent PG and PG-G production paths. (*A*) Dominant Reaction paths for PG Production Vary with AA and 2-AG Concentration. Each individual plot depicts the amount of flux through each path in 1*C* for a given concentration of 2-AG across varying concentrations of AA. Colors correspond to labeled paths in Fig. 1*B*. The error bars in each plot indicates the flux variation resulting from inferred kinetic rates. (*B*) Dominant Mechanisms of PG-G Production Vary with AA Concentration. Each individual plot is at a given concentration of 2-AG. In all plots AA increases from left to right at concentrations of 0.5, 1, 2, 4, 8 and 16 *µ*M in *A* and 0, 0.5, 1, 2, 4, 8, and 16 *µ*M in *B*. Colors correspond to labeled paths in Fig. 1*C* The error bars in each plot indicates the flux variation from inferred kinetic rates.

In the absence of 2-AG and at low (0.5 *µ*M) AA, PG is produced without allosteric modulation (Fig. 2*A*, purple; purple-labeled path in Figure 1C, top); as the concentration of AA increases, the proportion of PG produced with AA as an allosteric modulator also increases (Fig. 2*A*, green). When 2-AG is added to the system, PG production shifts to using 2-AG as an allosteric modulator (Fig. 2*A*, red), with this path favored to a greater extent as the concentration of 2-AG increases (Fig. 2*A*, lower plots). Even in the absence of 2-AG, about 20% of PG is produced by AA-modulated COX-2, and once even a small amount of 2-AG (0.5 *µ*M) is added to the system, more than half of PG production occurs via a 2-AG or AA allosterically modulated path. In the presence of high concentrations of either modulator, as much as 90% of PG is produced via an allosterically modulated path.

Because 2-AG and COX-2 display substrate-dependent inhibition (Mitchener et al., 2015), the production of PG-G occurs via fewer paths than are available to PG. In the absence of AA, all PG-G produced is generated in the absence of an allosteric modulator (Fig. 2B, purple), because the intermediate with 2-AG bound in both catalytic and allosteric sites is not turned over. As AA is added to the system, the proportion of PG-G produced by the AA-modulated pathway (Fig. 2*B*, red) increases. Thus, in the range of tested substrate concentrations, the dominant mechanism of PG-G production depends entirely on the amount of AA present in the system. Compared to PG, a smaller proportion of PG-G produced by the system results from an allosterically regulated pathway because PG-G is only created via the AA-modulated species or the allosterically unbound species. Nevertheless, at high concentrations of AA, again as much as 90% of PG-G is produced by AA-modulated COX-2. For paths containing a species bound in the allosteric site, binding at the catalytic site followed by binding at the allosteric site is the favored mechanism.

We note that at any given substrate concentration, the uncertainty arising from the calibrated kinetic parameter distributions never exceeds a 20% change in the percentage of product produced by a given path (Fig. S4-S8). We find that changes in substrate levels and their relative ratios have a much larger effect on the dominant reaction paths than changes in kinetic rates within the calibrated CORM parameter distributions. Overall, these findings suggest that variation of substrate concentrations in physiologically-relevant ranges has a significant impact on COX-2’s mechanism of catalysis.

### Pathway Entropy is Dynamic Across Input Concentrations

Calculating the flux through each path allows us to obtain information about the preferred sequences of reactions that the system executes while processing AA and 2-AG. However, these measurements do not provide an estimate of how chemical traffic (i.e. the flow of chemical signals in the network) is distributed throughout the network. To explore the distribution of biochemical network traffic, we introduce the *pathway entropy* to quantify the degree to which COX-2 utilizes multiple paths at different concentrations of substrates. Our definition of entropy, originally introduced by Claude Shannon (Shannon, 1948) provides a measure of the uncertainty in a probability distribution across states as follows:

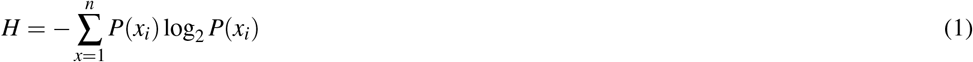

where *H* is entropy and *P*(*x*_*i*_) is the probability of any state *x*_*i*_. To determine the degree of uncertainty associated with product production (the pathway entropy), we considered each pathway as a state and use the fraction of flux that a given pathway contributes to the product as a measure for the probability of that state. This analysis yields a measure of how evenly distributed production is across possible paths. In general, evenly distributed fluxes across paths in a network would maximize pathway entropy for a multi-path system.

Since the dominant paths vary with substrate concentration (Fig. 2), we would expect that pathway entropy would also vary. In Fig. 3 we present the pathway entropy dependence on input concentration for PG (Fig. 3*A*) and PG-G (Fig. 3*B*). The pathway entropy for PG production is highest at intermediate levels of AA and low levels of 2-AG, while the pathway entropy is highest for PG-G production at intermediate levels of AA and any level of 2-AG. These maxima correspond to states where the reaction flux is most spread across the possible paths from reactant to product (see Fig. 2*A*, top plot, center, and Fig. 2*B*, top plot, center). In contrast, in the lowest entropy states - low AA and high 2-AG for PG (Fig. 2*A*, bottom plot, far left) and low AA across the entire 2-AG spectrum for PG-G (Fig. 2*B*, bottom row), flux is concentrated in a single or a few paths. Reaction flow is thus highly distributed in some conditions yet highly concentrated in one path in other conditions. This finding suggests that MIMO networks utilize multiple execution modes across input concentrations. It also suggests that approaches to modulate or inhibit network activity, which focus on disrupting one or more of these paths, may need to be tailored to specific conditions. These behaviors could have physiological relevance. For example, high-entropy conditions with highly redundant path fluxes may require multiple targets for inhibition compared to a condition with low entropy.

**Figure 3.**
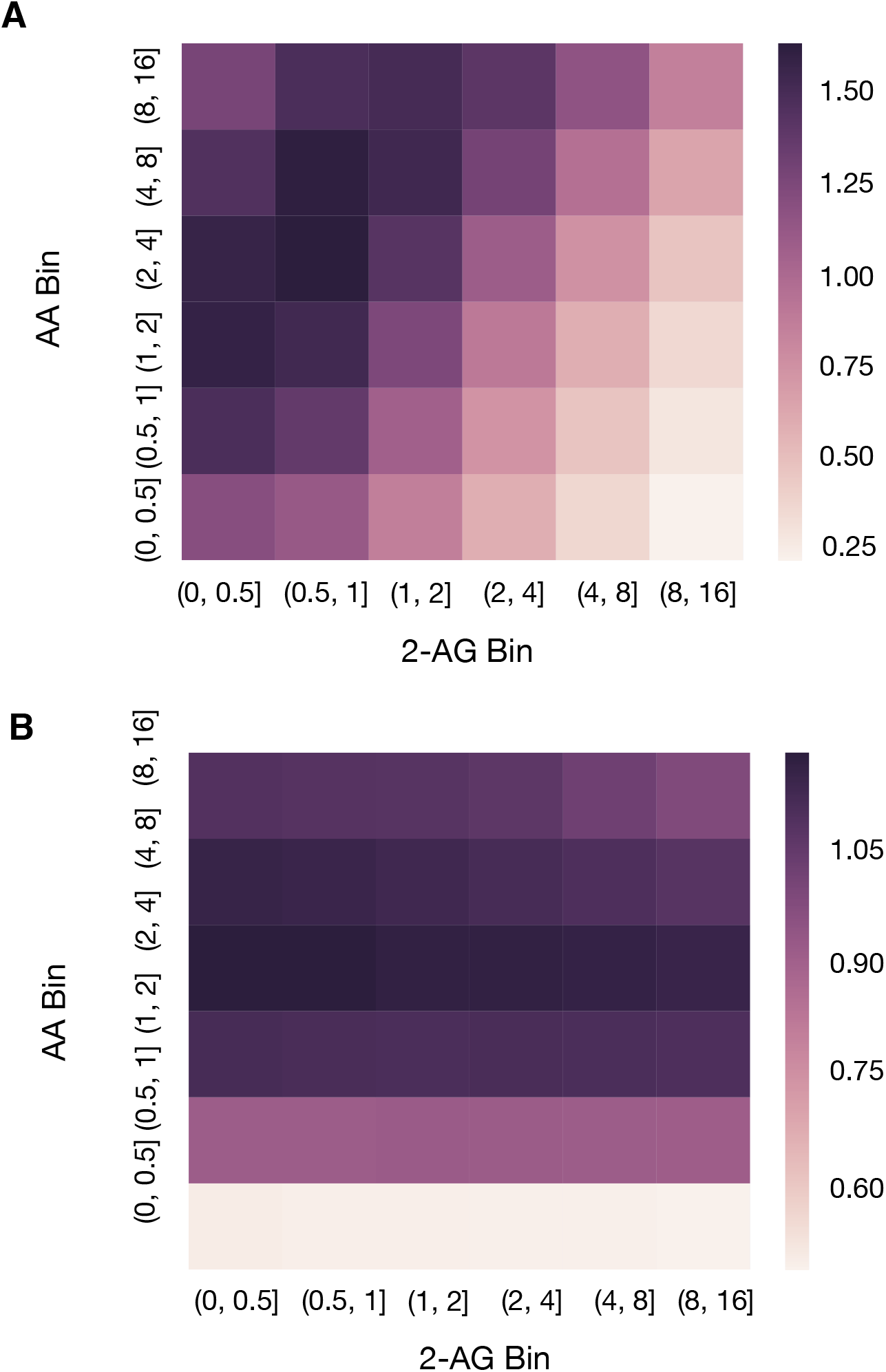
Pathway entropy within CORM. (*A*) Pathway Entropy for Production of PG. The intensity indicates the pathway entropy in units of bits. (*B*) Pathway Entropy for Production of PG-G. Units are the same as in *A*.

### Input Output Behavior in CORM

The above findings on pathway entropy suggest a complex relationship between input concentrations, reaction intermediates, and product concentration in CORM. To understand these relationships, we next considered concentration-dependence curves derived from simulations using a fixed set of CORM kinetic parameters in which PG was calculated at increasing AA concentrations in the presence of random quantities of 2-AG (Fig. 4*A*) or PG-G was calculated at increasing 2-AG concentrations in the presence of random quantities of AA (Fig. 4*B*). Each data point was taken at steady-state (10 seconds) for consistency with experiments and previous work. Note that the presence of both substrates results in competitive inhibition with suppression of product formation from either one. Thus, the highest levels of output in each case occur when the concentration of the opposing substrate is low. These levels are similar for PG and PG-G because COX-2 utilizes the two substrates with similar catalytic efficiencies when they are present individually. As the concentration of the opposing substrate increases, competitive inhibition is partially balanced by positive allosteric modulation in the case of the conversion of AA to PG, but exacerbated by negative allosteric modulation in the case of the conversion of 2-AG to PG-G. Therefore, the suppression of PG-G formation is greater than that of AA formation as seen in the lower plateau level achieved in (Fig. 4*B*). In addition, the range of inputs over which the output varies depends significantly on which input/output pair is chosen (note the difference in that range in Fig. 4*A,B*). Clearly, variation of *both* inputs (e.g. changing AA *in addition* to changing 2-AG in Fig. 4*A*), results in significant variation in the outputs. Thus, while our simulations are deterministic, introducing uncertainty in the AA concentration generates a type of “extrinsic noise” in the relationship between 2-AG and PG-G (Fig. 4*B*), and *vice versa* for the impact of 2-AG on the relationship between AA and PG, Fig. 4*A*). This noise represents allosteric modulation in the network due to varying input concentrations.

**Figure 4.**
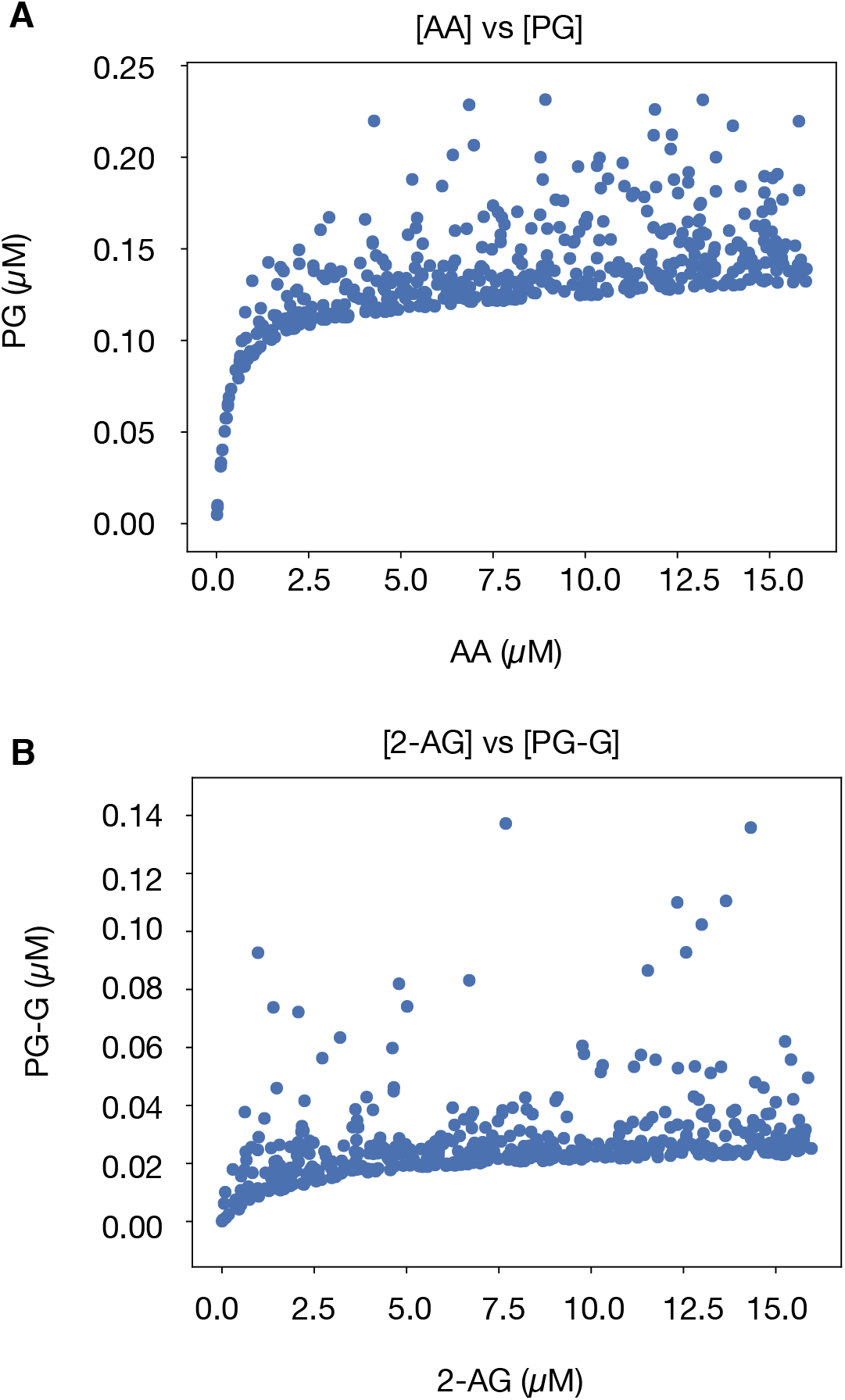
Input vs output plots for substrates and products in CORM. (*A*) Input vs Output plots for AA to PG. 2-AG varies randomly. All concentrations are measured at steady-state (10 seconds). (*B*) Input vs Output plots for 2-AG to PG-G. AA varies randomly. All concentrations are measured at steady-state (10 seconds).

### Channel Capacity from Substrates to Products

To better understand how this output variation, combined with the shape of the concentration-dependence curves, influences the COX-2 reaction network, we applied an additional concept from information theory to measure dependence between inputs and outputs, namely the Mutual Information:

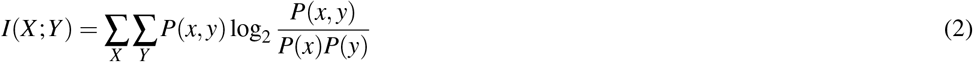

where *X* represents a given signal and *Y* the response to that signal (Shannon, 1948). Mutual information quantifies the degree to which one variable provides information about a second variable. Equivalently, it is a measure of how knowledge about one variable decreases uncertainty in the value of a second variable. For biological systems, quantifying mutual information is challenging because the input distribution is generally unknown. Previous work (Cheong et al., 2011; Selimkhanov et al., 2014; Suderman et al., 2017) has focused on estimating the “channel capacity,” which is the maximum information attainable across all possible input distributions:

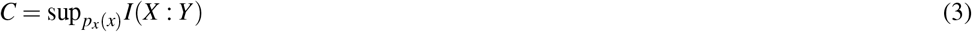

Note that any practical calculation provides a lower bound estimate for the channel capacity *C*, since only a finite set of input distributions is used to estimate *I* (Suderman et al., 2017). We calculated channel capacities using the approach and software published in Suderman *et al.* (Suderman et al., 2017), which is similar to that used in Cheong *et al.* (Cheong et al., 2011).

We applied this estimate to two different sets of simulations. In the first set of simulations, we considered a case where AA and 2-AG are perfectly correlated with each other; to do this, we sampled the AA concentration from a uniform distribution on [0, 16 *µM*] and set the 2-AG concentration to be exactly the same. In the second set, we *independently* sampled the input AA and 2-AG substrate concentrations from a uniform distribution on the interval [0, 16 *µM*]. In each case, we sampled a total of 500 distinct input conditions and ran CORM simulations to 10s to agree with experiments and previous work (Mitchener et al., 2015). The channel capacity was then estimated between the two different inputs (either AA or 2-AG) and every possible intermediate and product. The maximum theoretical channel capacity, log_2_(500) ≈ 9 bits, would be obtained if each of the 500 inputs yielded a distinct response. We repeated the channel capacity calculation for the top 5000 most probable parameter vectors from the calibrated parameter ensemble. This then allowed us to quantify the effect of kinetic parameter variation on channel capacities in the system. In total the analysis required approximately 1.5M CPU hours. An example of input data used for calculating channel capacities from AA to PG and 2-AG to PG-G for a single parameter set is shown in Fig. 4. Greater detail is provided in the SI Methods.

### COX-2 Integrates Information from Both AA and 2-AG

For ease of visualization, we estimated kernel densities of channel capacities given variation in calibrated kinetic parameters as shown in the violin plots in Fig. 5. In these plots, the data are represented by a central box plot that provides the mean, interquartile range and 95% credible interval, and the surrounding shape depicts the probability distribution, with wider regions indicating a higher probability. Because the input-output relationship in these simulations is deterministic, deviations from the theoretical maximum (≈ 9 bits) arise from the two phenomena described above: either changes in the input do not really lead to significant changes in the output (*i.e.* the “flat” part of the concentration-dependence curves in Fig. 4) or the independent variation in one of the substrates generates variation in the output that is not due to the input being considered (*i.e.* the apparent noise in Fig. 4).

**Figure 5.**
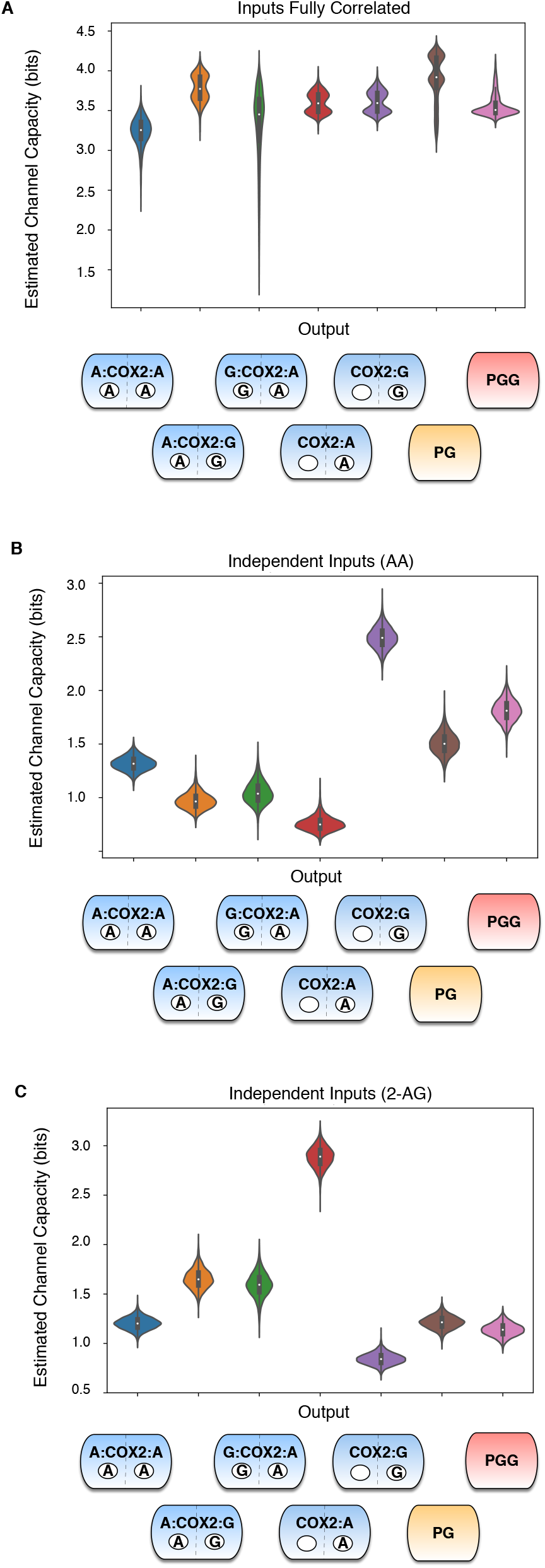
Estimated channel capacities from substrates to intermediates or products in CORM. (*A*) Estimated Channel Capacities from Input to intermediates and final products within CORM when levels of AA and 2-AG are strongly correlated (Pearson correlation coefficient = 1). Distributions in the channel capacities arise from uncertainty in the kinetic parameter values after model calibration.(*B*) Estimated Channel Capacities from AA to intermediates and final products within CORM when AA and 2-AG are varied independently. Distributions in the channel capacities arise from uncertainty in the kinetic parameter values after model calibration.(*C*) Estimated Channel Capacities from 2-AG to intermediates and final products within CORM when AA and 2-AG are varied independently. Distributions in the channel capacities arise from uncertainty in the kinetic parameter values after model calibration.

From Fig. 5, it is clear that the combination of these effects significantly reduces the observed channel capacities from the theoretical maximum. The highest observed value for any of the input/output pairs (AA to PG, 2-AG to PG-G, etc.) is at most half of the theoretical maximum (less than 4.5 bits). When input values are perfectly correlated ([AA] = [2-AG]), Fig. 5*A*, the channel capacity between the (correlated) inputs and the outputs is between 3 and 4.5 bits (depending on the parameters), indicating that, while not perfect, the concentration-dependence curves allow for high levels of information flow between inputs and outputs. It is interesting to note that the uncertainty in the kinetic parameters leads to some variation in the calculated channel capacities; since the inputs here are correlated, this variation is due to changes in the shape of the concentration-dependence curves between data sets. Many channel capacities in the correlated case are bimodal, suggesting that two specific concentration-dependent curve shapes are most likely.

When the inputs are varied independently, channel capacity values decrease even further (Fig. 5*B* and *C*). The channel capacity between AA and PG or PG-G is generally less than 2 bits, and the channel capacity between 2-AG and those outputs is generally less than 1.5 bits. This could occur for two reasons. First, a lack of correlation could result in less entropy in the response (*i.e.* less uncertainty in the value of the product). Since the mutual information is limited by the response entropy (eq. 2, (Shannon, 1948; Cheong et al., 2011; Suderman et al., 2017)), this would cause a decrease in the mutual information. However, if the response entropy remains constant when there is no correlation between inputs, then mutual information can only decrease if information transfer through the network is less efficient. As shown in Fig. S9, the response entropy does not differ between the independent and correlated cases, indicating that independent variation in one of the inputs while the other input is known has a large effect on the output. In other words, COX-2 is truly an integrator of these signals, since accurate determination of the substrate concentrations given the output is considerably more difficult if the two substrates are independently varied.

Since perfect correlation and complete independence represent only the two extremes of the relationship between AA and 2-AG concentration, we also investigated the behavior of the system when the inputs exhibit moderate correlation (Pearson correlation coefficient = 0.5), and when the inputs are consistently present in a 2-to-1 AA-to-2-AG ratio (Fig. S10 and Fig. S11). The behavior when input ratios were fixed was similar to that for the correlated values (when the input levels were fixed equal to each other); channel capacities were again higher than in the independent case and the effect of kinetic parameter variation on channel capacity was higher. When the inputs are moderately correlated, the system is still able to obtain high channel capacities for some kinetic parameter sets, although the overall distribution of channel capacities shifts to lower values compared to when input correlation is perfect, further confirming COX-2 input integration.

### Information Flow is Dictated by Substrate Concentration

We next tested whether the channel capacity between substrates and products varies with substrate level. We binned the input data into four quadrants (high or low values of either substrate) and calculated the channel capacity between inputs and outputs independently for each quadrant; input ranges were otherwise identical to those used for the calculations described above. Low substrate values spanned 0-8 *µ*M and high substrate values 8-16*µ*M. Both independently varied inputs (Fig. 6*A*) and correlated inputs (Fig. 6*B*) yielded estimated channel capacities that were significantly different between the different regions of input space. In addition to differences in PG and PG-G channel capacity, we found that the distribution of information that passed through different intermediates changed with substrate concentration (Fig. S13 and Fig. S14); certain paths to product had greater information transfer capacity at particular levels of substrates. This echoes findings from our pathway analysis (Figs. 2 and 3), indicating that changes in substrate concentration result in significant changes in how the enzyme executes its catalytic mechanism. Interestingly, we found no detectable correlation between the flux through a pathway and the mutual information between an input and an intermediate in that path (Fig. S15 and Fig. S16). We leave further investigation of the relationship between information transfer and actual physical reaction fluxes for future work.

**Figure 6.**
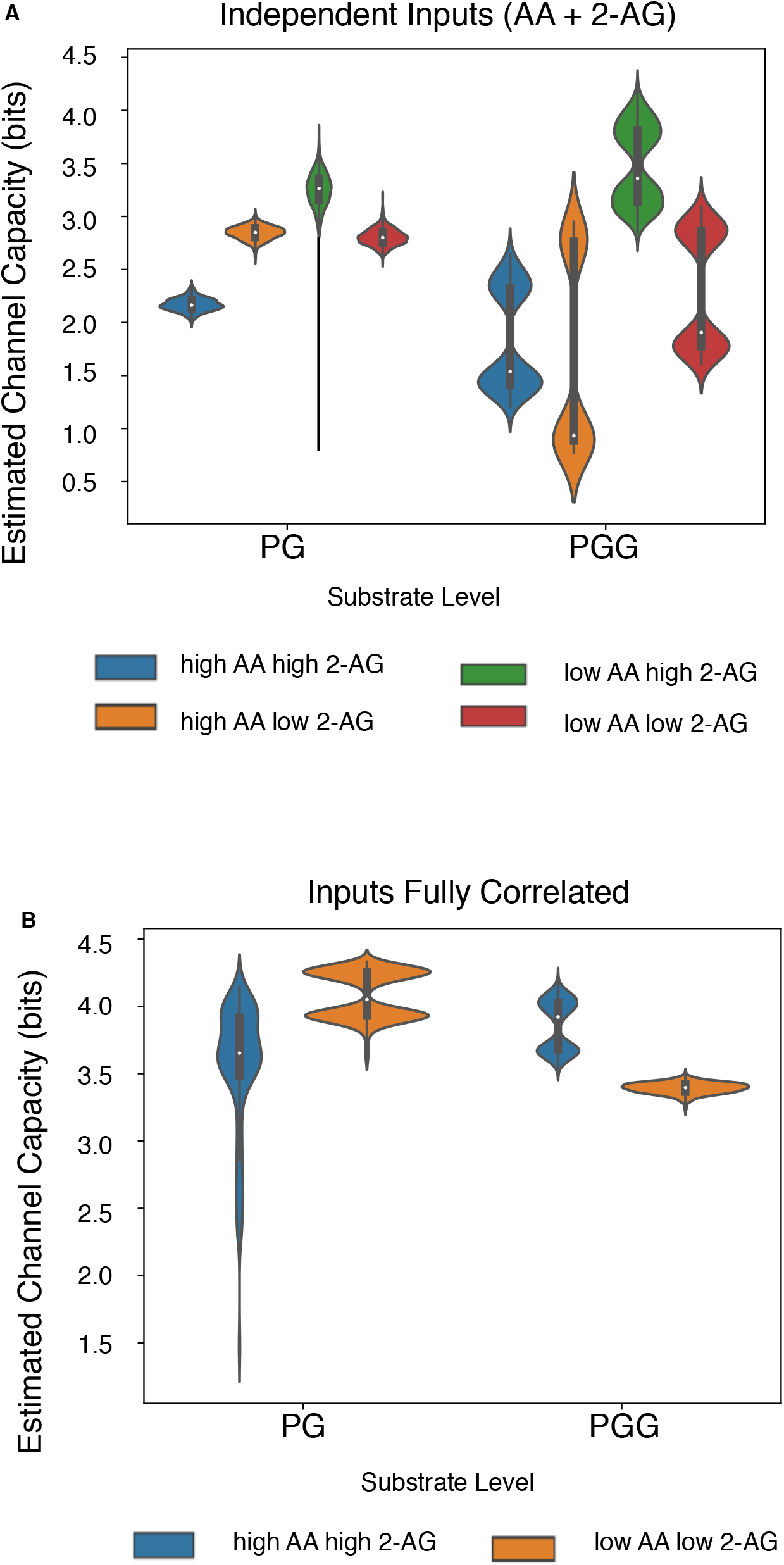
Effect of substrate level on estimated channel capacities between substrates and products in CORM. (*A*) Total Estimated Channel Capacity from AA and 2-AG combined to products across regions of substrate space. Distributions in the channel capacities arise from uncertainty in the kinetic parameter values after model calibration. (*B*) Estimated Channel Capacities from input to products when levels of AA and 2-AG are perfectly correlated across regions of substrate space. Distributions in the channel capacities arise from uncertainty in the kinetic parameter values after model calibration.

Splitting the input space into different quadrants also revealed signficant variation between different parameter sets, with most distributions showing significant bimodality across parameters (Fig. 6). This suggests that both the shape of the concentration-dependence curves, and the impact of “extrinsic noise” due to variation of one substrate independent of another, varies across parameter sets. Since all of these parameter sets are equally consistent with experimental data (Mitchener et al., 2015), this suggests that multiple modes of information flow are available to the COX-2 reaction network without significant changes to the core functionality of the enzyme.

## DISCUSSION

*In vivo*, COX-2, AA, and 2-AG concentrations vary across cells in different tissues (Seibert et al., 1997; Monjazeb, 2006; Sugiura et al., 2006). In most tissues, AA processed by COX-2 is released from membrane phospholipids, predominantly through the action of cytosolic phospholipase A2 (Leslie, 2015). In some tissues, (particularly the brain) a major source of AA is hydrolysis of 2-AG (Ignatowska-Jankowska et al., 2014; Long et al., 2008). In turn, 2-AG is also sourced from membrane phospholipids; through the sequential action of phospholipase C, which forms diacylglycerol (DAG), followed by conversion of DAG to 2-AG by DAG lipase (Fezza et al., 2014). Both DAG lipase and cytosolic phospholipase A2 are stimulated by increases in intracellular Ca^2+^ (Leslie, 2015; Bisogno et al., 2003). Thus, many stimuli (such as zymosan phagocytosis by macrophages (Rouzer and Marnett, 2005)) promote the release of AA and 2-AG simultaneously, with concentrations of AA typically higher than those of 2-AG. Considering the precursor-product relationship between 2-AG and AA, however, it is conceivable that in some cells, the levels of the two substrates may change inversely to one another, or that the level of one may change while the other remains constant. These considerations suggest that the system features we find that vary with AA and 2-AG level (pathway entropy and information transfer capacity) are states accessible by the true biological system with the attendant repercussions for information transfer within that system. In addition, the postulated link between diet and the substrates available for COX-2 turnover (Chen, 2010) suggests that the information transfer properties of the system could be modulated by fatty acid intake.

COX-2 has significant regulatory flexibility: it is an allosteric protein, with multiple substrates and multiple allosteric regulators, all of which can influence how COX-2 operates on its substrates *in vivo*. The pathway analysis (Fig. 1*B* and 1*C*) suggests that COX-2 functions by first binding a substrate at the catalytic site, followed by binding of an allosteric regulator. Allostery can be viewed as a shift in the conformational free-energy landscape sampled by COX-2 through preferential binding of the allosteric regulator to particular conformations (Lechtenberg et al., 2012; Nussinov and Tsai, 2013). From this perspective, modulating the concentrations of allosteric regulators in the COX-2 system shifts the conformational ensemble towards conformations favored by particular regulators. In the case of PG, these conformations are more easily turned over to product than the unmodulated enzyme, while for PG-G, the allosteric influence makes catalysis less energetically favorable (shifts the ensemble towards conformations that are less active). This allows COX-2 to manage the balance between PG and PG-G production in a more complex (and potentially farther-reaching) fashion than that provided by simple competition between substrates. This added complexity suggests a physiological reason why the COX-2 system would integrate information from multiple inputs: by adding a second competitive input, the system can access different responses than with a single input. Furthermore, the response dynamics of COX-2 gain even greater complexity because its inputs act as allosteric modulators in addition to substrates. The situation *in vivo* is likely far more complicated (and flexible) than considered here, as COX-2 has potential substrates in addition to AA and 2-AG (Rouzer and Marnett, 2009), and some nonsubstrate fatty acids that act as allosteric regulators (Dong et al., 2016b; Yuan et al., 2009; Dong et al., 2011, 2013, 2016a). In addition, many of the non-steroidal anti-inflammatory drugs that target COX-2 also may bind at either the catalytic or allosteric site.

One advantage of this complexity may be the significant robustness of this system to variation in the kinetic parameters. The 5000 parameter sets we considered here all fit experimental data on PG and PG-G production equally well, despite variation of over three orders of magnitude in some of the parameter values (Mitchener et al., 2015). Our results on both pathway flux (Fig. 2) and information flow (Fig. 5) indicate that different parameter sets favor different distributions of paths from substrate to product, and transfer information through the network in different ways. Yet the overall function of the enzyme is the same despite all of this variation. *In vivo*, a change in the kinetic rates could correspond to a mutation or a change in the level of molecular crowding for the reaction. The availability of multiple “modes of execution” in this complex enzyme thus allow the system to be highly robust to such changes. This complex architecture could also allow the system to be highly evolveable through a mechanism of facilitated variation (Gerhart and Kirschner, 2007). We expect that future work on parameter variation will reveal major insights into the evolution of robustness in enzymes like COX-2.

In this work we applied a systems biochemistry framework to understand chemical reaction flux, pathway entropy, and information flow in the COX-2 system and investigate how these adjust to dynamic input concentrations and correlations. The methods and approach utilized here could be applied to further probe the COX-2 system by including more inputs (its other substrates, allosteric regulators, and inhibitors), or transferred to a larger, more complex network. Given the complexity present in even the simple network considered here, we predict that a systems biochemistry approach to larger networks would provide non-intuitive insights into the dynamics of the system as a whole.

## METHODS

### Modeling and Model Calibration

CORM was encoded as a PySB (Lopez et al., 2013) model containing 13 distinct biochemical species and 29 chemical reactions. It was calibrated to experimental data consisting of PG and PG-G concentrations at steady state across a range of substrate concentrations (Mitchener et al., 2015; Shockley et al., 2017).

## Calculating Path Fluxes and Channel Capacities

The method for determining paths of production and the total flux through a path is described in detail in SI Methods and Fig. S1. Channel capacities were calculated using the method from (Cheong et al., 2011) and the software of (Suderman et al., 2017). Extended detail is available in SI Methods.

The datasets generated during and/or analysed during the current study are available from the corresponding author on reasonable request.

## ACKNOWLEDGMENTS

We would like to thank Dr. Blake A. Wilson for critical reading of the manuscript. We gratefully acknowledge funding from the National Science Foundation (1411482 to C.F.L.), National Cancer Institute (U01CA215845 to V.Q. and C.F.L.), Defense Advanced Research Projects Agency (Cooperative Agreement no. W911 NF-14-2-0022 to C.F.L.).

## AUTHOR CONTRIBUTIONS

Conceptualization: EMS, CFL; Methodology: EMS, CFL, EJD; Investigation: EMS, CFL, EJD, CAR, LJM; Resources: CFL, LJM; Analysis: CFL, EMS, EJD, CAR; Writing original draft: EMS, CFL; Writing Review and Editing: EMS, CFL, EJD, CAR, LJM; Supervision: CFL; Funding Acquisition: CFL, LJM

