## Supplemental Text for "Signal integration and information transfer in an allosterically regulated network"

### Methods

#### *Model Simulations*

All CORM simulations were run using the PySB Lopez et al. (2013) SciPy-based integrator. PySB version 1.1 (available at http://github.com/LoLab-VU/pysb) and CORM version 1.0 (available at http://github.com/LoLab-VU/CORM) were used. A detailed description of the CORM is available in the supplementary material in Shockley et al. (2017) For flux analysis, species and reaction trajectories were generated at twelve time points between .01 and ten seconds. Calibrated parameter values from previous work Mitchener et al. (2015) were used. Credible intervals for the ensemble of parameter values (taken from Mitchener et al. (2015)) are shown in Table 1. Catalytic rates are in units of $s^{-1}$ and disassociation constants in units of $\mu$M.

#### *Pathway Analysis*

If one considers the network of biochemical interactions included in CORM, there are a multitude of paths from enzyme and substrates to either possible product. To enumerate these paths, we employ a graph-theory based approach. In this approach, a node edge graph is constructed in which each node is a set of reactants or products, and each edge represents a reaction connecting a reactant-product set. The edges are directed with the direction dictated by the net integrated reaction flux ($k_f - k_r$) at a given time; these edges are therefore time dependent given the time dependence of the net reaction flux. We then determine all simple paths (paths with no repeating reactant-product nodes) connecting initial reactants (COX-2 + AA or COX-2 + 2-AG in this case) to final products (PG and PGG) via the net integrated flux directed edges. The relationship between paths then allow us to calculate the proportion of flux each path contributes to the final product as the joint probability of all reactions that constitute that path (see Fig. 1).

#### *Channel Capacity Calculations*

Channel capacities were estimated using the EstCC package introduced in Suderman et al. (2017), which calculates mutual information between a channel input and output using the method of Cheong et al. (2011). EstCC input data was comprised of the results of CORM simulations at a given calibrated parameter vector for 500 different initial AA values drawn from M $U(0, 16)$, after a ten second simulation. For independent input values, 2-AG values were also drawn from M $U(0, 16)$. For strongly correlated inputs, 2-AG values were set to the value of AA. AA or 2-AG was considered the signal and the concentration of intermediate or final PG/PGG product the response. The number of signal bins and the number of response bins was selected with EstCC using the procedure described in Suderman et al. (2017). The estimate of channel capacity for a given input (and input correlation) and output was performed separately for simulations with 5000 different calibrated parameter sets to allow estimation of the uncertainty in channel capacity stemming from parameter uncertainty.

For the channel capacity calculations in a given region of substrate space, calculations were performed identically to the original analysis, except that input values were constrained to low (0-8 M) or high (8-16 M) levels of AA and 2-AG, and only 500 calibrated parameter sets were used to estimate uncertainty from parameter uncertainty (to make the calculations computationally feasible).

In total all channel capacity calculations required approximately 1.5 million CPU hours to complete. Computation was performed using a dynamically scaled Amazon Web Services (AWS) Batch cluster containing up to 20,000 CPUs.


[H]

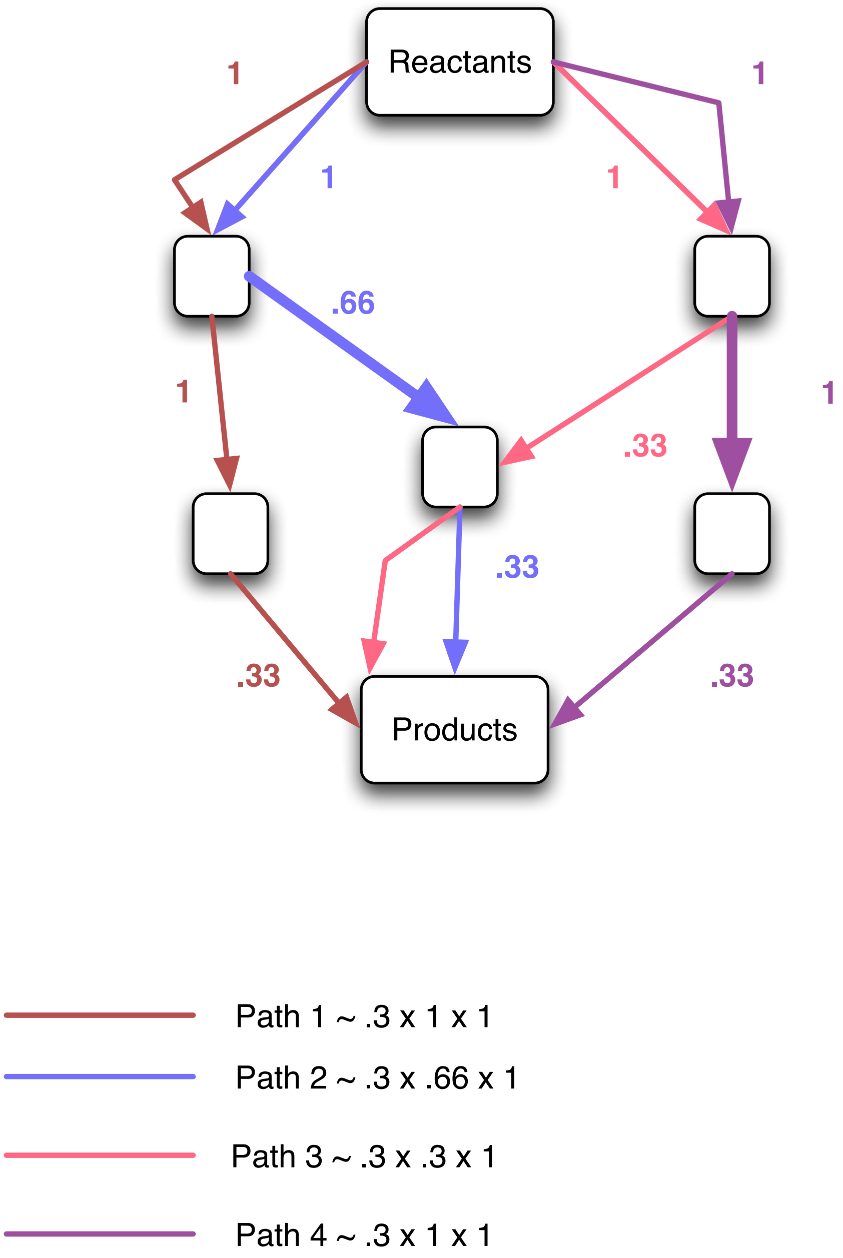


**Figure 1.** Calculating paths between reactants and products. Edges are directed towards the flow of the net integrated reaction flux (for bidirectional reactions) or the integrated forward reaction flux (for unidirectional reactions). Edge labels indicate the probability of selecting that reaction path into a node, calculated as the proportion of total flux leaving the node that came from a given reaction.

| Parameter | Lower Bound | Upper Bound |
|---|---|---|
| kcat_AA2 | 0.92 | 17 |
| kcat_AA3 | 1.3 | 1.7 |
| kcat_AG3 | 0.14 | 0.73 |
| KD_AA_allo1 | 4.9 | 3700 |
| KD_AA_allo2 | 1.2 | 2.9 |
| KD_AA_allo3 | 0.20 | 1.2 |
| KD_AA_cat2 | 0.0 | 0.060 |
| KD_AA_cat3 | 0.0 | 0.24 |
| KD_AG_allo1 | 13 | 5400 |
| KD_AG_allo2 | 0.23 | 4.6 |
| KD_AG_cat2 | 0.010 | 3.9 |
| KD_AG_cat3 | 0.0 | 0.070 |

**Table 1.** Credible Intervals for Parameters in CORM

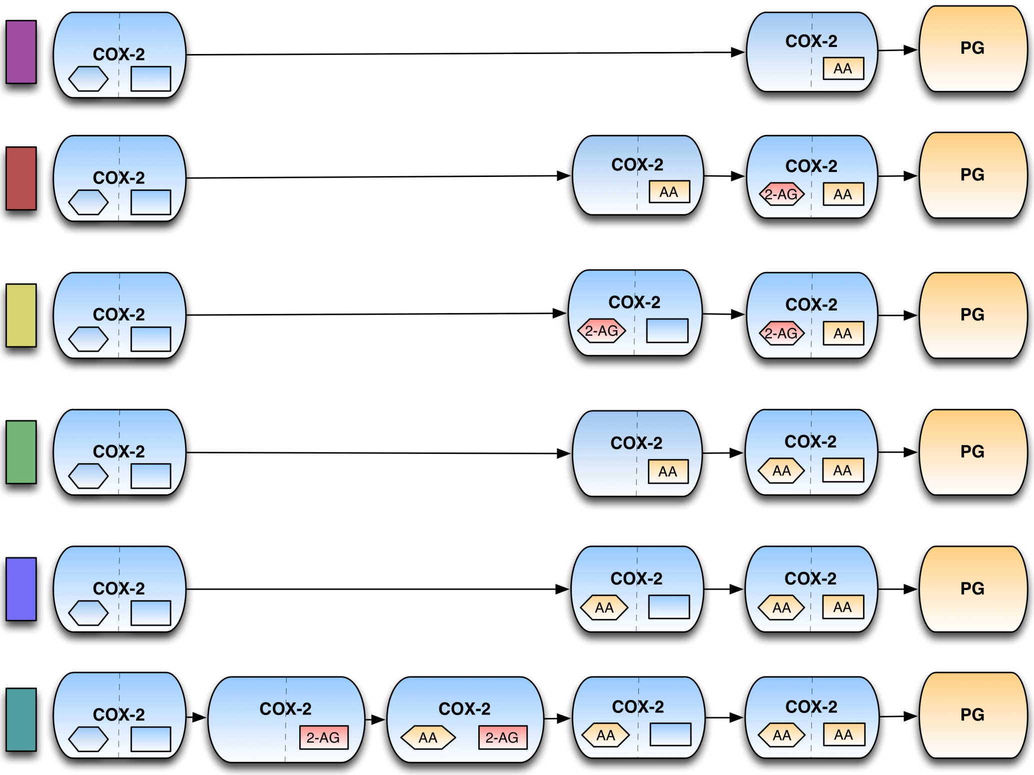


**Figure 2.** Possible paths between COX-2 and PG in CORM.

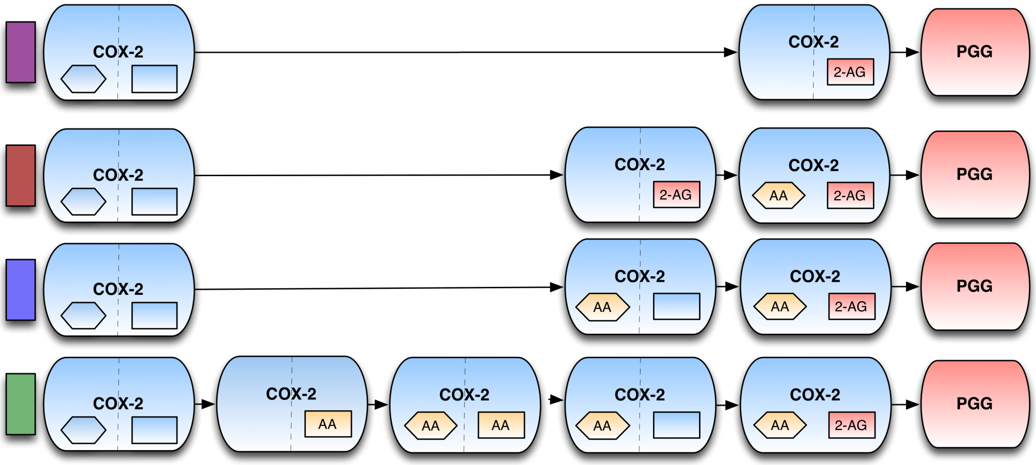


**Figure 3.** Possible paths from COX-2 to PGG in CORM.

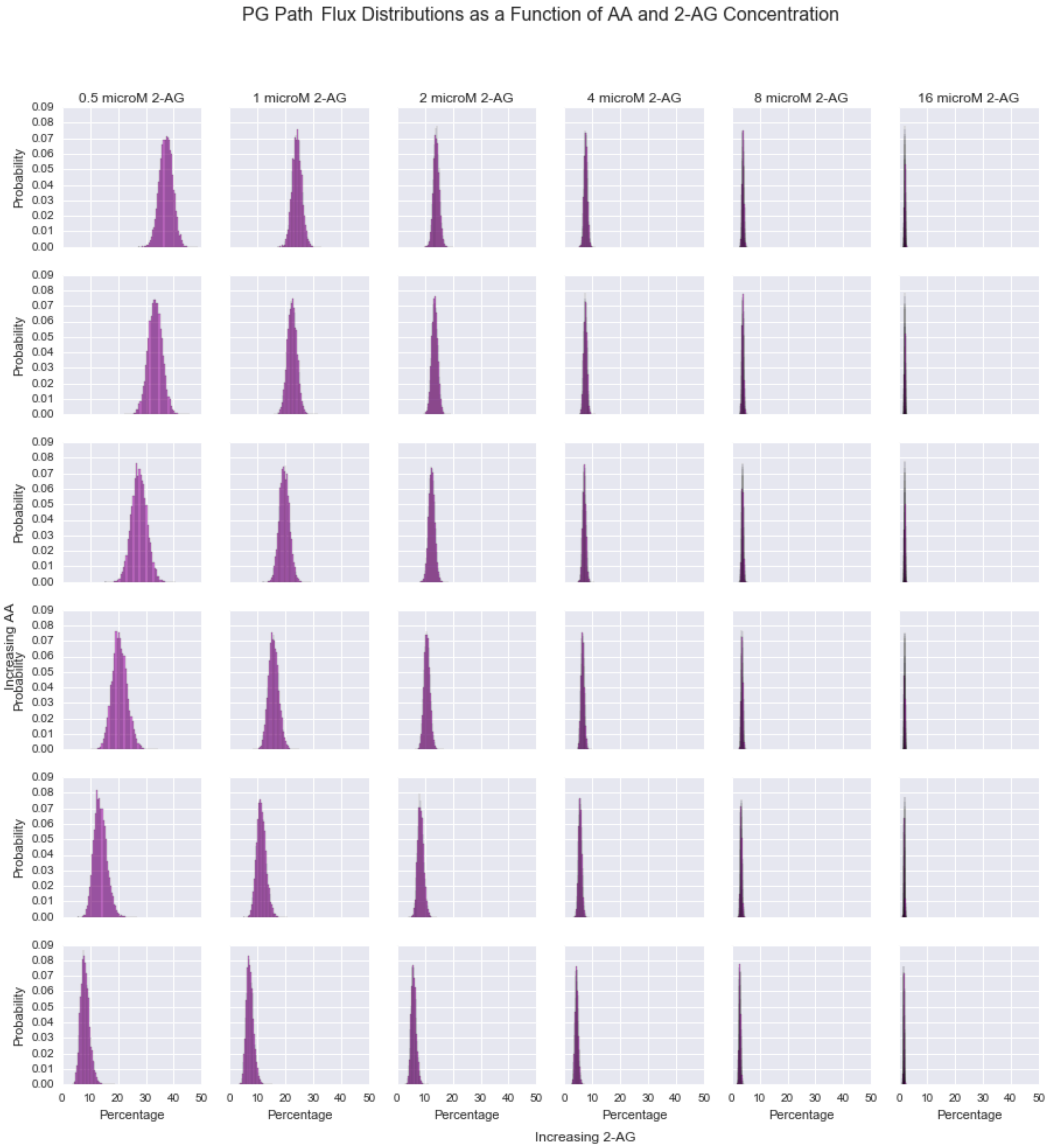


**Figure 4.** Distribution (arising from calibrated parameter uncertainty) of the percentage of the total flux to product PG from the path COX-2 ¡-¿ COX-2:AA -¿ PG at different starting AA and 2-AG concentrations.

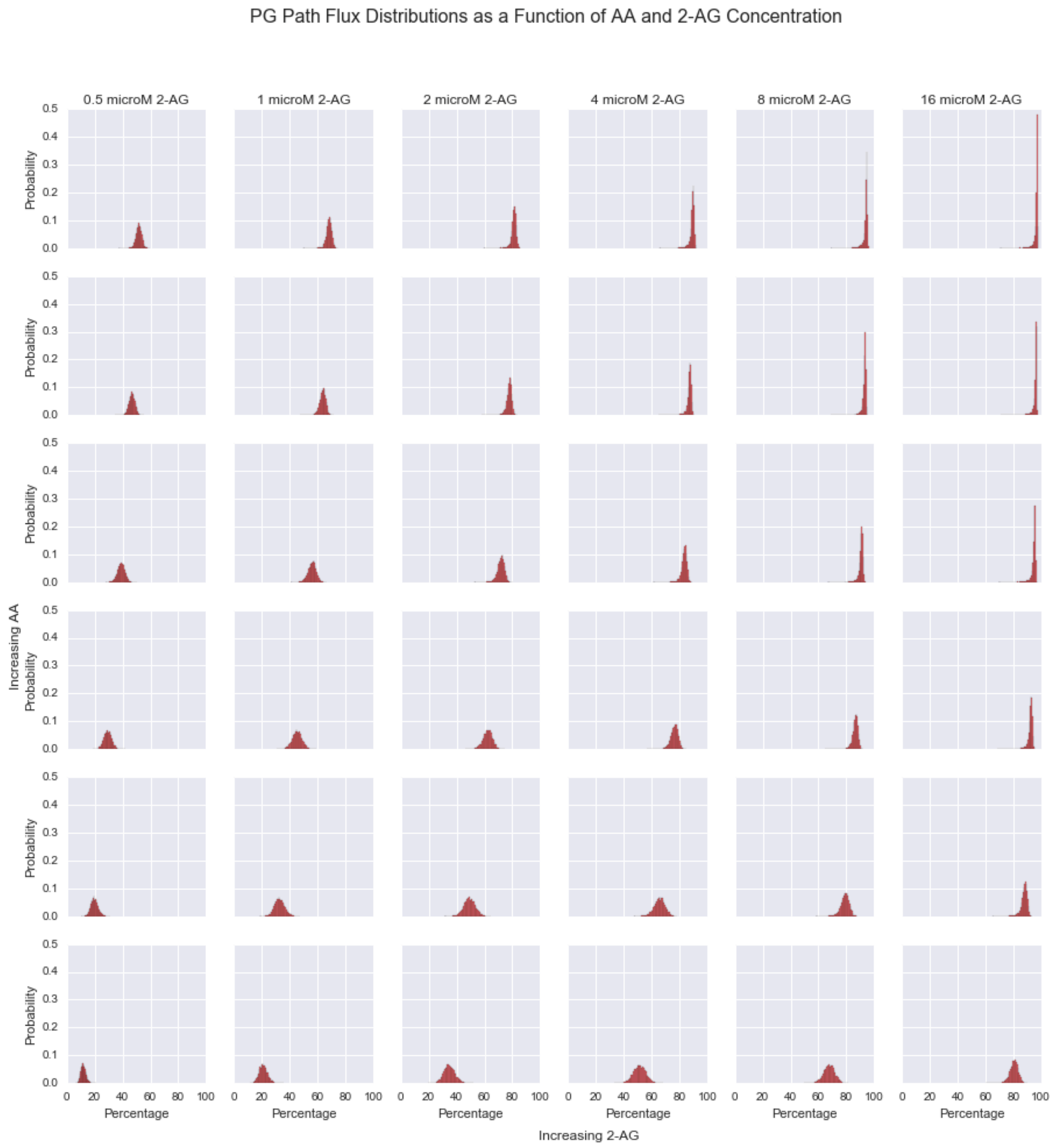


**Figure 5.** Distribution (arising from calibrated parameter uncertainty) of the percentage of the total flux to product PG from the path COX-2 ¡-¿ COX-2:AA ¡-¿ 2-AG:COX-2:AA -¿ PG at different starting AA and 2-AG concentrations.

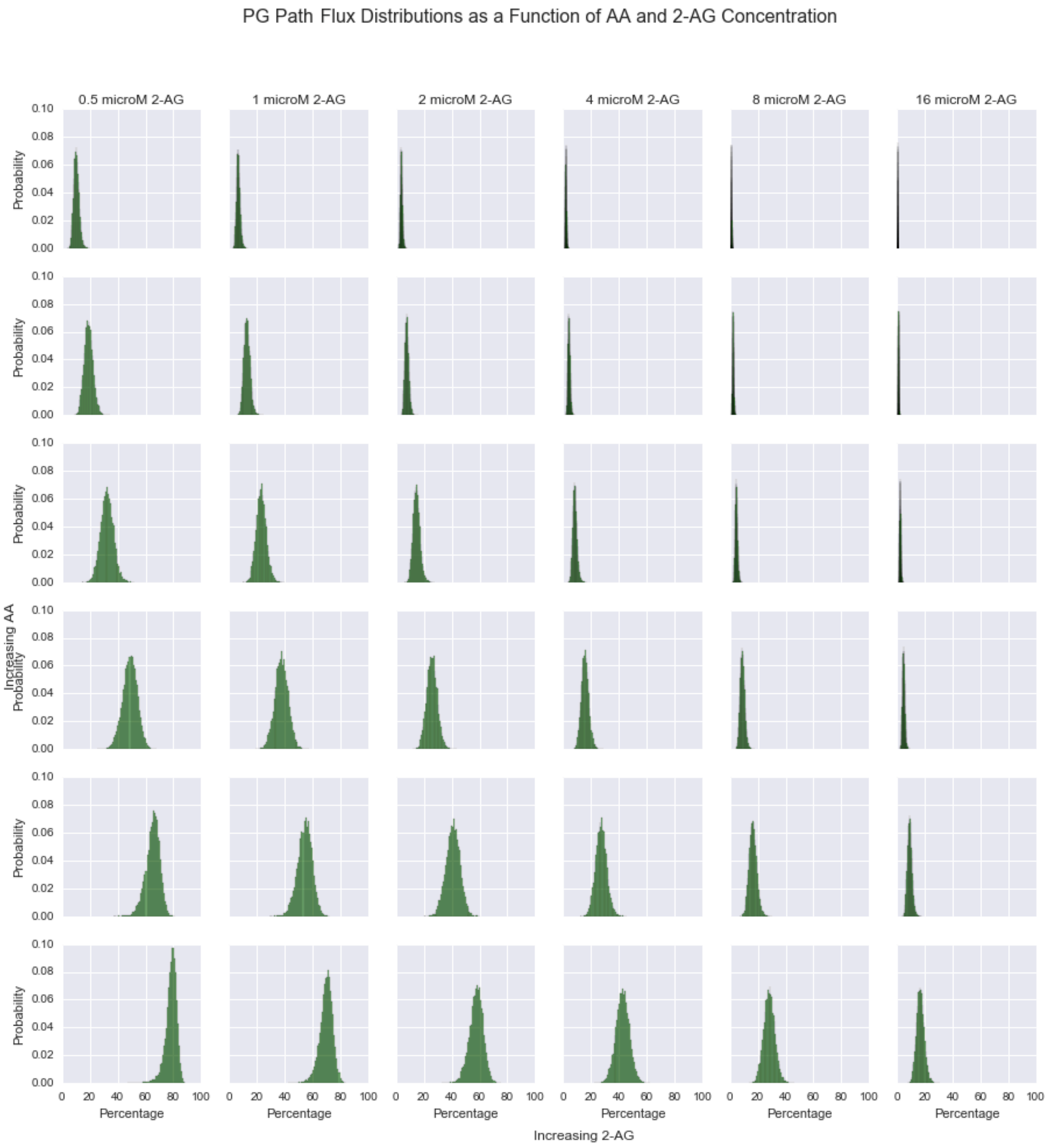


**Figure 6.** Distribution (arising from calibrated parameter uncertainty) of the percentage of the total flux to product PG from the path COX-2 ¡-¿ COX-2:AA ¡-¿ AA:COX-2:AA -¿ PG at different starting AA and 2-AG concentrations.

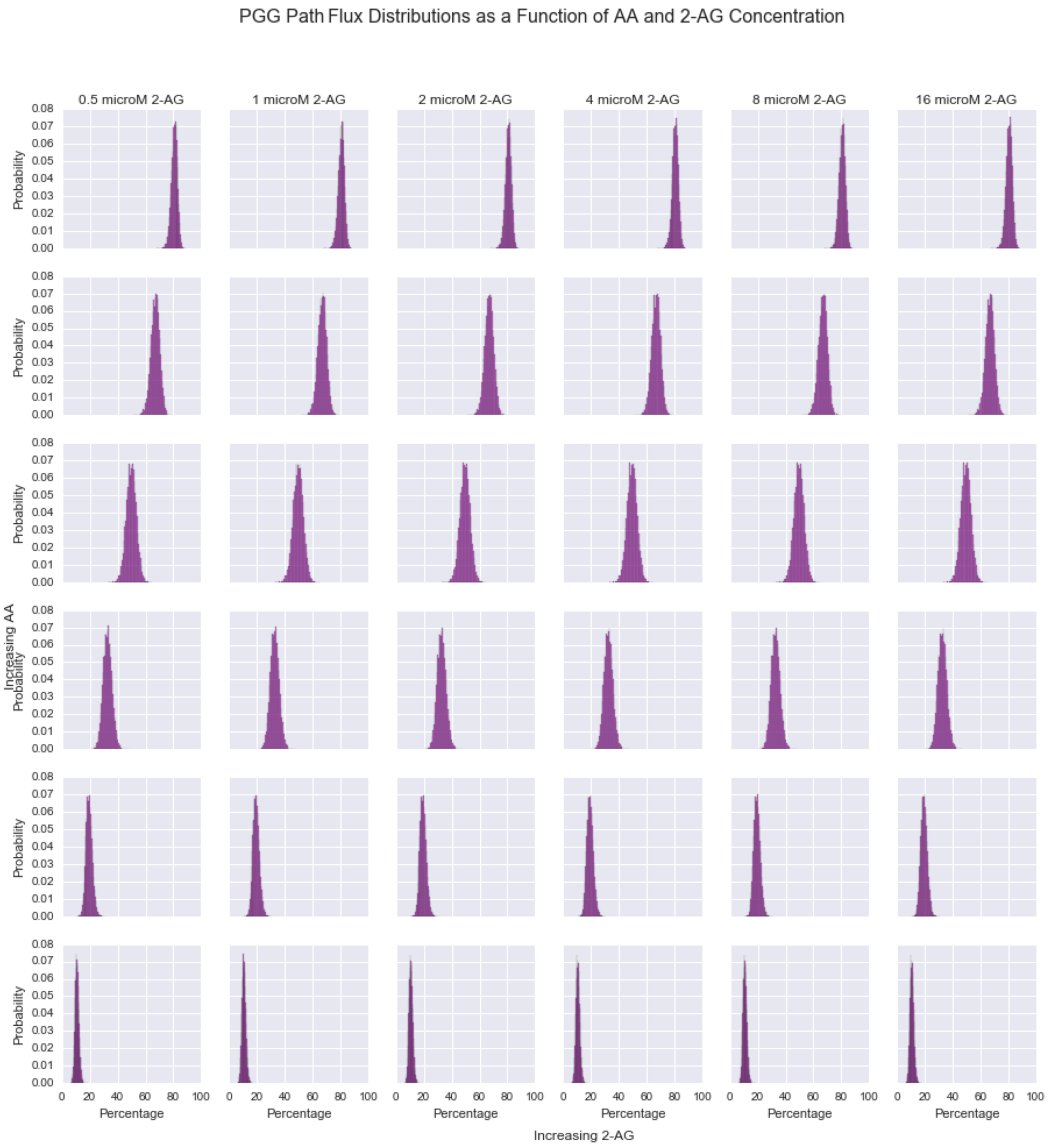


**Figure 7.** Distribution (arising from calibrated parameter uncertainty) of the percentage of the total flux to product PGG from the path COX-2 ¡-¿ COX-2:2-AG -¿ PGG at different starting AA and 2-AG concentrations.

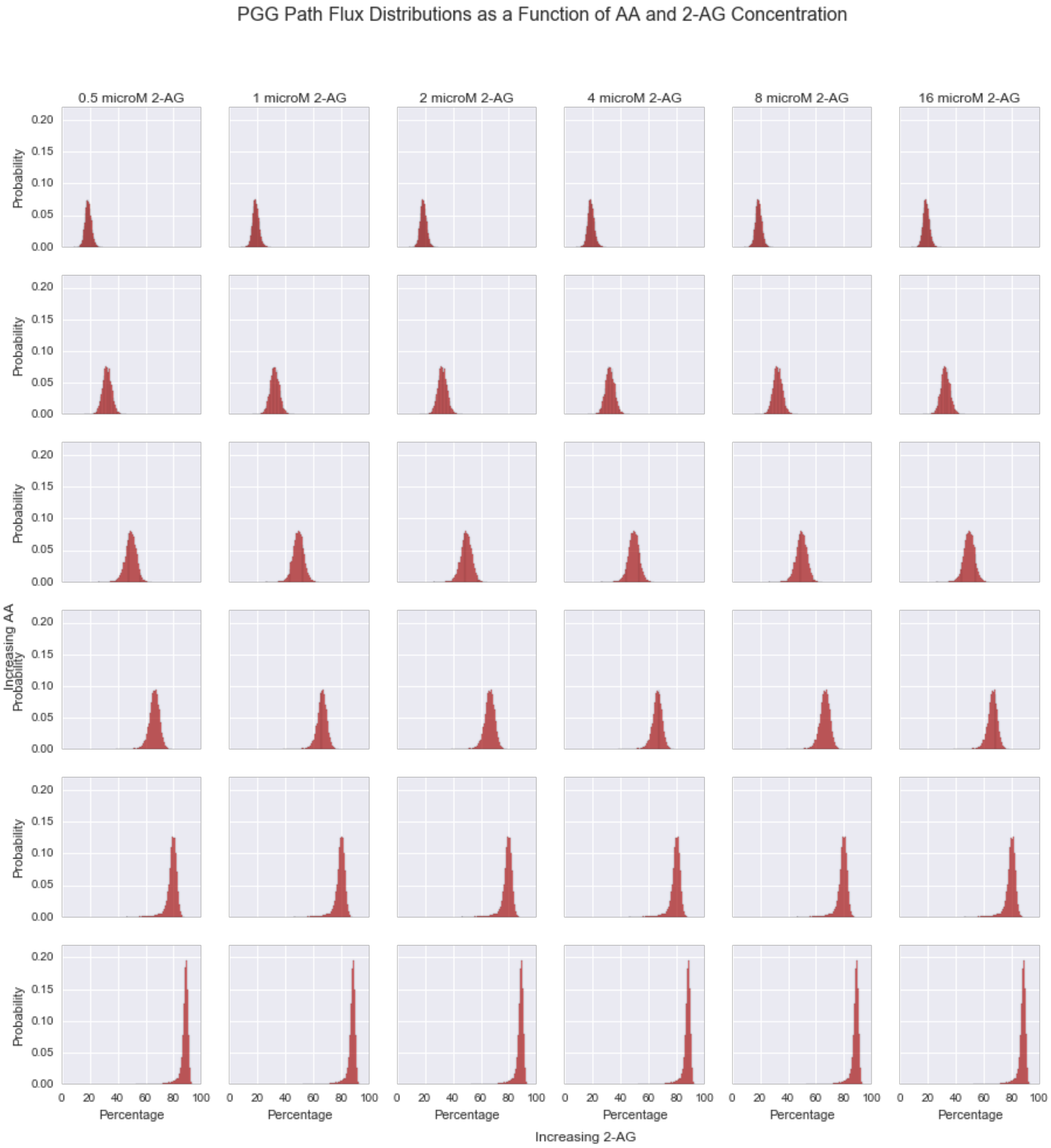


**Figure 8.** Distribution (arising from calibrated parameter uncertainty) of the percentage of the total flux to product PGG from the path COX-2 ¡-¿ COX-2:2-AG ¡-¿ AA:COX-2:2-AG -¿ PGG at different starting AA and 2-AG concentrations.

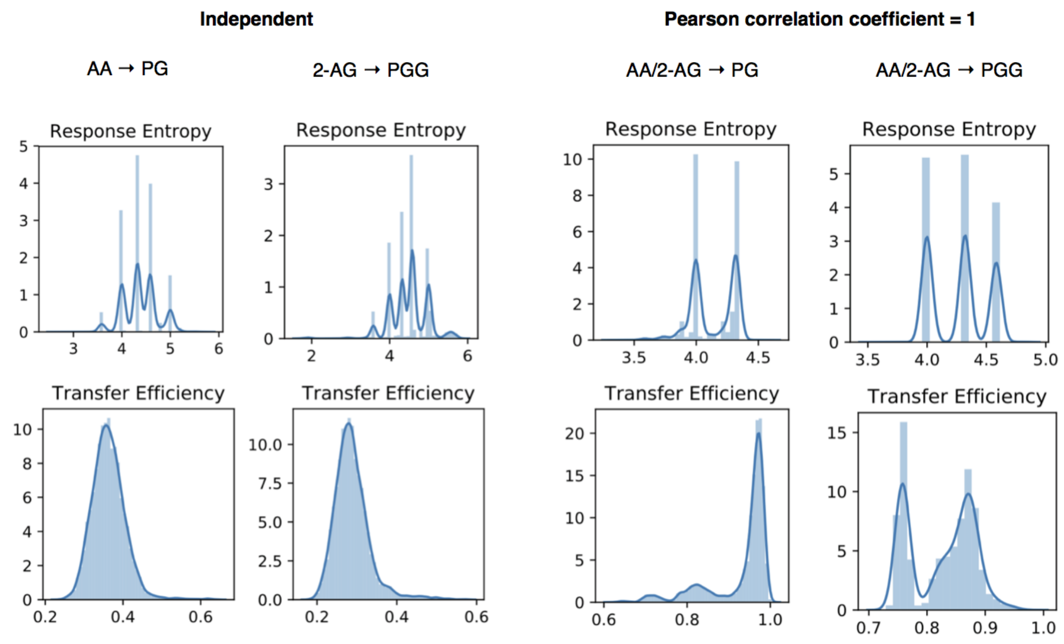


**Figure 9.** Distributions of response entropy under different input correlation and signal/response pairs. While the response entropy spans a similar range with both independent and correlated inputs, the transfer efficiency increases with correlation.

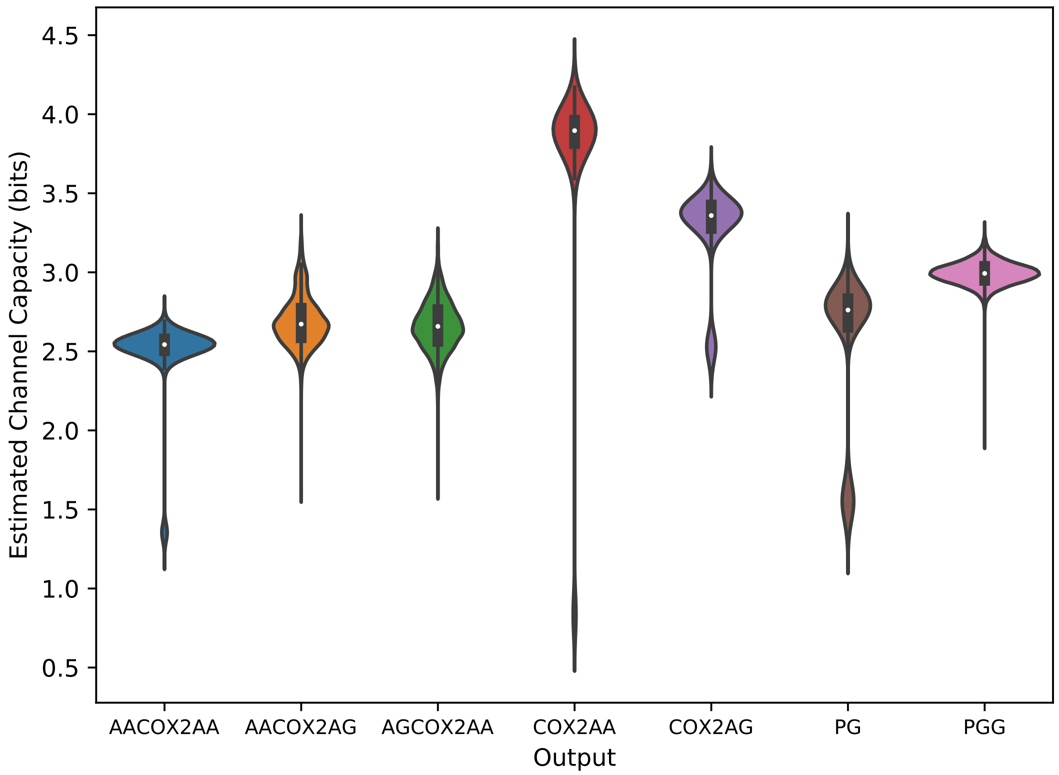


**Figure 10.** Sum of channel capacity from AA and 2-AG to intermediates and final outputs when inputs are semi-correlated (Pearson correlation coefficient = .5). Distributions arise from uncertainty in calibrated parameter values.

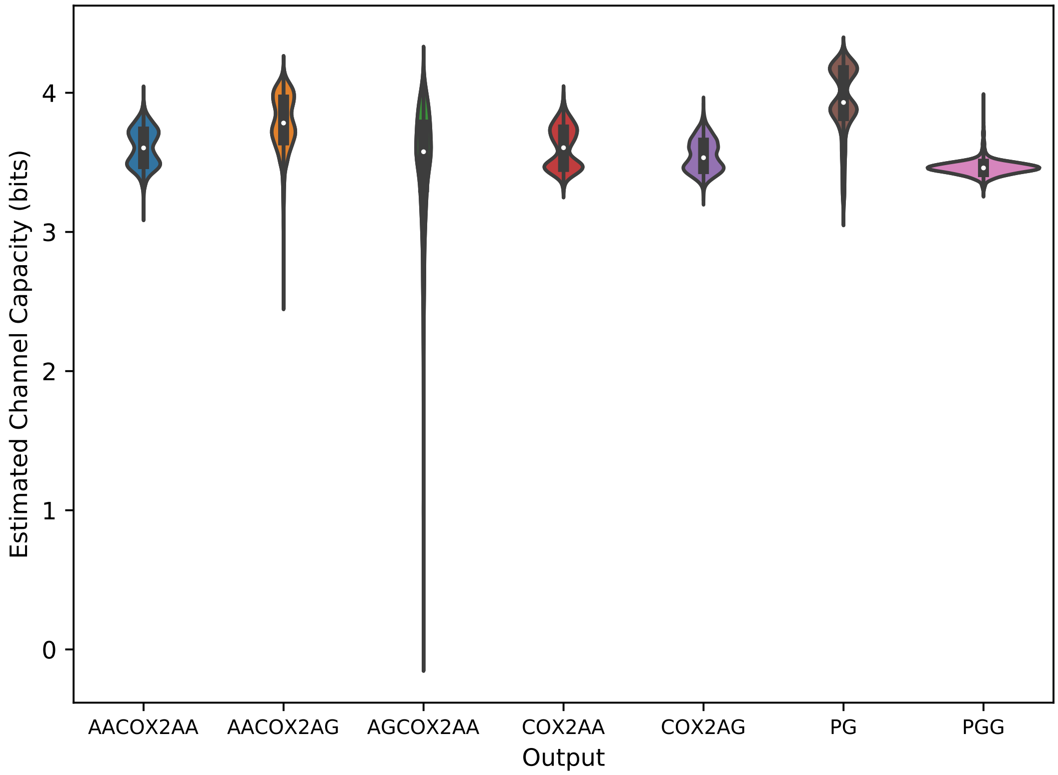


**Figure 11.** Channel capacity from AA and 2-AG to intermediates and final outputs when inputs are present in a 2 to 1 AA to 2-AG ratio. Distributions arise from uncertainty in calibrated parameter values.

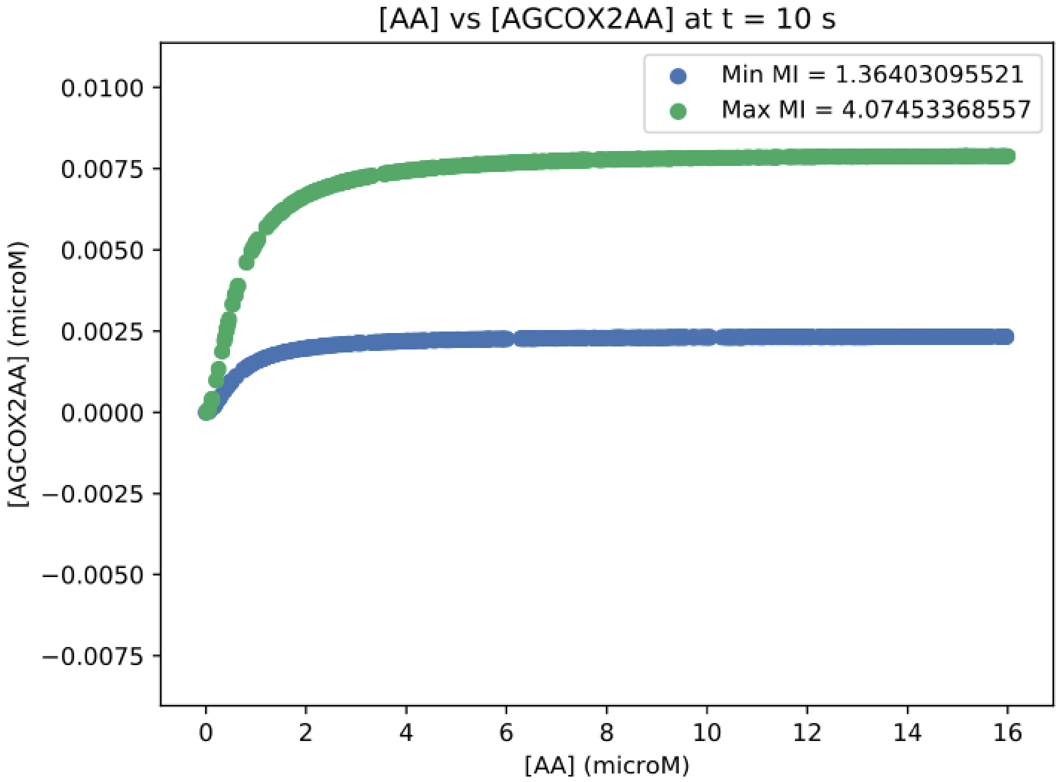


**Figure 12.** Input-Output response for 2-AG:COX-2:AA for parameter sets yielding maximum and minimum channel capacitys.

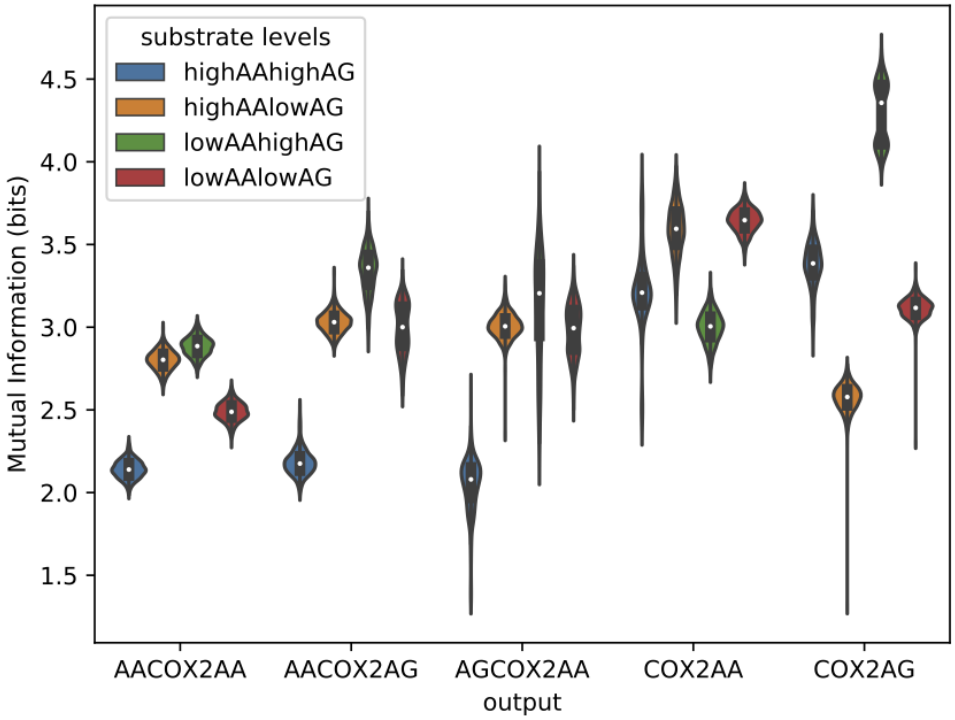


**Figure 13.** Sum of channel capacity from AA and 2-AG to intermediates when inputs are varied independently in different regions of substrate space. Distributions arise from uncertainty in calibrated parameter values.

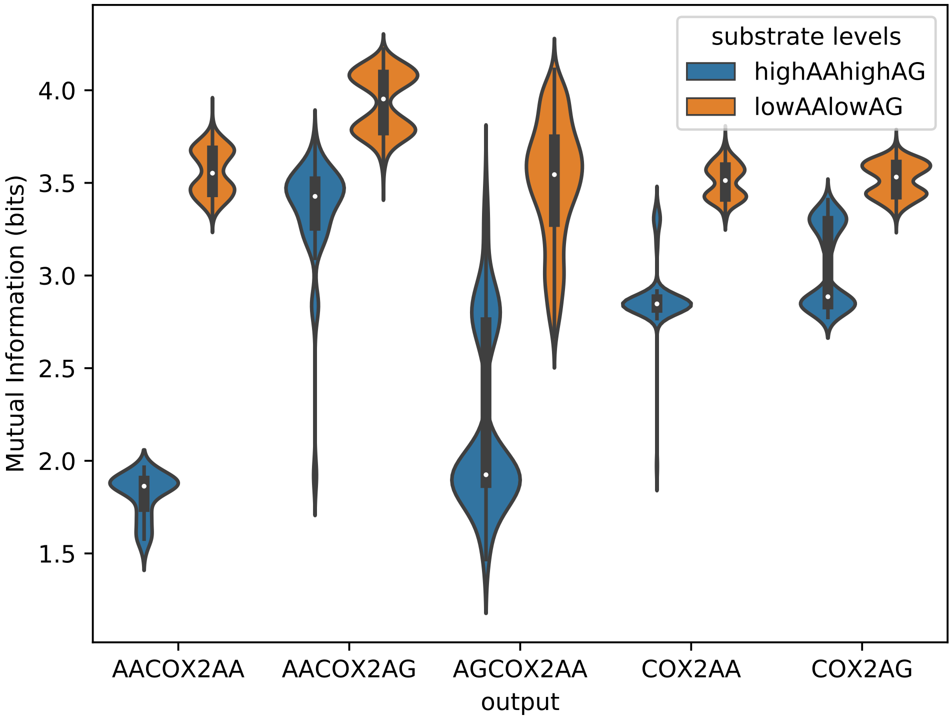


**Figure 14.** Channel capacity from AA and 2-AG to intermediates when inputs are varied correlated (Pearson correlation coefficient = 1) in different regions of substrate space. Distributions arise from uncertainty in calibrated parameter values.

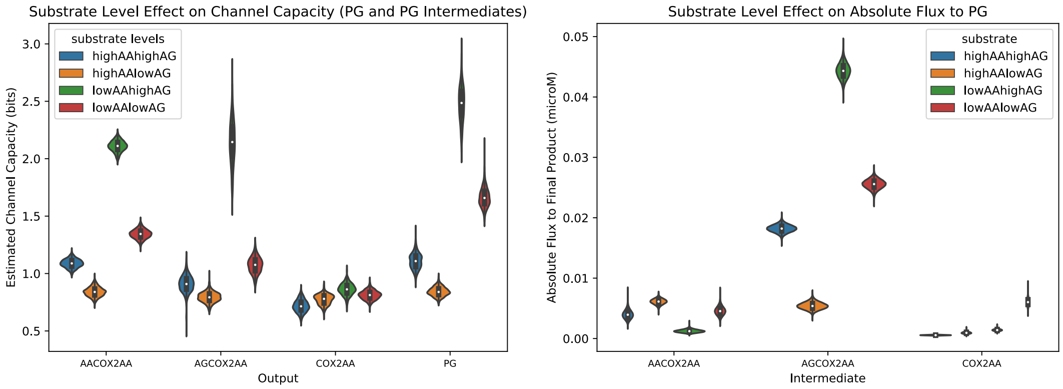


**Figure 15.** Left: channel capacity between independent inputs and PG intermediates at different substrate levels. Right: absolute flux between independent inputs and PG intermediates at different substrate levels.

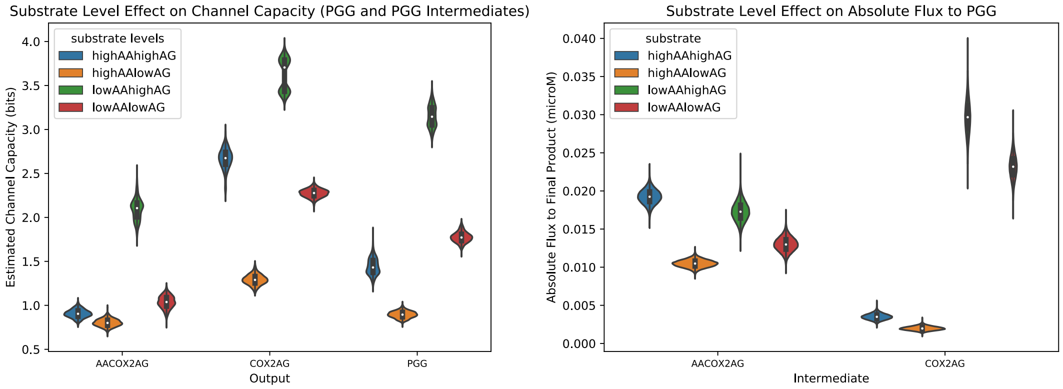


**Figure 16.** Left: channel capacity between independent inputs and PGG intermediates at different substrate levels. Right: absolute flux between independent inputs and PGG intermediates at different substrate levels.
